# An alignment free approach confirms semantic properties of species proteomes

**DOI:** 10.1101/2021.06.04.447028

**Authors:** Antonio Starcevic, Ena Melvan, Toni Cvrljak, Janko Diminic, Jurica Zucko, Paul F. Long

**Author notes:** Correspondence to: Antonio Starcevic. These authors contributed equally to this work (Ena Melvan, Toni Cvrljak).

## Abstract

Alignment-based methods dominate molecular biology. However, by primarily allowing one-to-one comparisons, these methods are focused on a gene-centered viewpoint and lack the broad context essential to analyze how complex biological systems function and evolve. In actuality, a gene is part of genome where more than one sequence contributes to the functional network and evolutionary trajectory of the cell. The need for conservation of established interactions, is arguably more important to the evolutionary success of species than conservation of individual function. To test whether such contextual information exists, a distributional semantics method - Latent Semantic Analysis (LSA), was applied to thousands of species proteomes. Using natural language processing, Latent Taxonomic Signatures (LTSs) were identified that outperformed existing alignment-based BLAST methods when random protein sequences were being mapped to annotated taxonomy according to GenBank. LTSs are a novel proteome distributed feature, suggesting the existence of evolutionary constraints imposed on individual proteins by their proteome context. Even orphan proteins are exhibiting LTSs, which makes their uniqueness linked to a specific taxonomic level questionable. Unlike more simple bias, LTSs represent a self-similarity pattern, where random sets of species proteins show the same statistical properties of a complete proteome at many scales. Natural language processing and machine learning provide insights not easily discernable using alignment based methods suggestive there is more to species related differences than just translational optimization.

## Background

The major role of DNA is information storage. This information enables function, development, growth and reproduction of all known organisms and is organized in the form of genes, gene clusters, chromosomes and ultimately genomes. If DNA represents the code of life, the way in which it is organized within cells as a genome, can be compared to the organization of a computer database or a structured document. Even the challenges modern databases face, such as data redundancy, archiving, versioning, replication etc. can be compared to genomes, with evolution constantly performing unscheduled updates. Using this analogy, the atomic information unit is the gene. A gene is defined as the basic physical and functional unit of heredity acting on stored information to make proteins, which in return perform actual work ensuring that the gene will multiply and persist through replication. Genomes, however, are composed of thousands of genes that participate in complex interaction networks. These networks must be conserved across evolutionary time, which poses a legitimate question as to how individual genes and proteins retain information related to inclusion in already established networks. Molecular evolutionary studies rely exclusively on alignment-based homology to infer relatedness, yet these methods often struggle to infer common biological function or interactions between sequences that lack sufficient homology to be aligned [1]. Latent Semantic Analysis (LSA) is a method based on natural language processing that can uncover information common to sequences sharing a network or a context but which share little or no homology [2]. The concept behind LSA is the Distributional hypothesis [3], whereby linguistic items with similar distributions have similar meanings. LSA has been extensively used in text processing applications e.g. analyzing relationships between sets of documents [4]. The same principle has been applied to studying evolution by analyzing the distribution of the 4-letter nucleotide code of genes, or the 20-letter amino acid one in proteins. Predating alignment-based methods, an observation that genomes have species-specific preference for nucleotides, summarized by Chargaff’s 2^nd^ rule [5] helped solve the double helical structure of DNA. Even the most prominent alignment-based methods, do not adhere completely to the concept of optimal alignment. Instead, these methods find k-mers, which are substrings of length *k,* used as a “signature” for the underlying sequence [6]. This heuristic approach has been applied in domain of genome assembly, where k-mers are used during the construction of De Bruijn graphs [7,8]. Despite the fact that k-mers do deviate from an ideal alignment concept, this approach has allowed for some of the major advances in molecular biology. Despite this, the majority of biologists remain sceptic of alignment-free approaches that teach away from established molecular dogma that similar sequences share similar function or structure, which is conserved through evolution if the sequences confer biological fitness[9]. Although not yet being fully convincing for the mainstream scientific majority, LSA has been successfully used to predict enzyme substrate specificity [10] and a variant of LSA called probabilistic latent semantic analysis (PLSA) was used to analyze proteomes of closely related pathogens [11]. Others have gone further and proposed bio-vectors referring to biological sequences in general and more specifically, protein-vectors for proteins and gene-vectors for DNA sequences. These vectors achieved >90% family classification accuracy, outperforming existing classification methods [12]. Herein, all available protein sequences from NCBI “nr” database were grouped according to source organism. Resulting species proteomes were used to create species LSA model that were correlated with the NCBI Taxonomy database in order to test whether reductionist approach favored by alignment-based methods producing a gene-centered view can be complemented by more integral, natural language processing method putting emphasis on context based information. The results indicated that there is more than just bias behind differences in species sequences. Instead of simple bias, actual self-similar signatures emerged, which were present in all of species proteins, including the ones currently being labeled as taxonomically restricted. In order to increase awareness of the broader audience alignment-free methods truly deserve, we made the LSA species model freely available for all to test on http://matrix.pbf.hr/ and we plan to update it on regular basis.

## Results

### Alignment-free approach to inferring species similarity

To assess whether different proteins making up a species proteome store context-based information, a taxonomy-benchmarking test was performed to compare the results with the NCBI Taxonomy database. Latent Semantic Analysis is able to utilize entire protein sets for both query and model creation. For this purpose, three protein query sets of different sizes were selected and deliberately removed from the remaining proteomes used for the LSA model. To ensure the same benchmarking conditions, initial removal of 500 proteins from each of the 54,526 proteomes provided the largest training set from which two smaller training sets were extracted (100 and 50 proteins each). The remaining proteome was used to build the LSA model in all three benchmarking tests. The model was built by transforming protein sequences into lists of “words”, in this case 3-peptide motifs based on a sliding window approach (Supplementary Figure 1A). These 3-peptide frequencies were then weighted, embedded in the form of species vectors and used for pairwise comparison based on cosine similarity (Supplementary Figure 1B). The taxonomy related prediction capacity of the LSA model was assessed by challenging the model with the three query sets of different size for each of the 54,526 species, using single best-hit approach (SBH) and a voting scheme method (VSM) (Supplementary Figure 1B). Briefly, for each taxon query sequence set, the SBH approach assigned taxonomy of the most similar subject taxa vector (from LSA training set), whilst the VSM method took the five most similar taxa vectors from the training set and used these in a voting scenario (Supplementary Figure 1C). The results using both SBH and VSM approaches gave significant agreement with the NCBI taxonomy, ranging from 66.9% - 78.9% for SBH and from 82.6% - 92% for VSM. The average taxonomy correlation calculated across all taxonomic ranks from the smallest (50 proteins) to largest (500 proteins) sample used in benchmarking tests are shown in Fig. 1, A, B and C.

**Fig.1.**
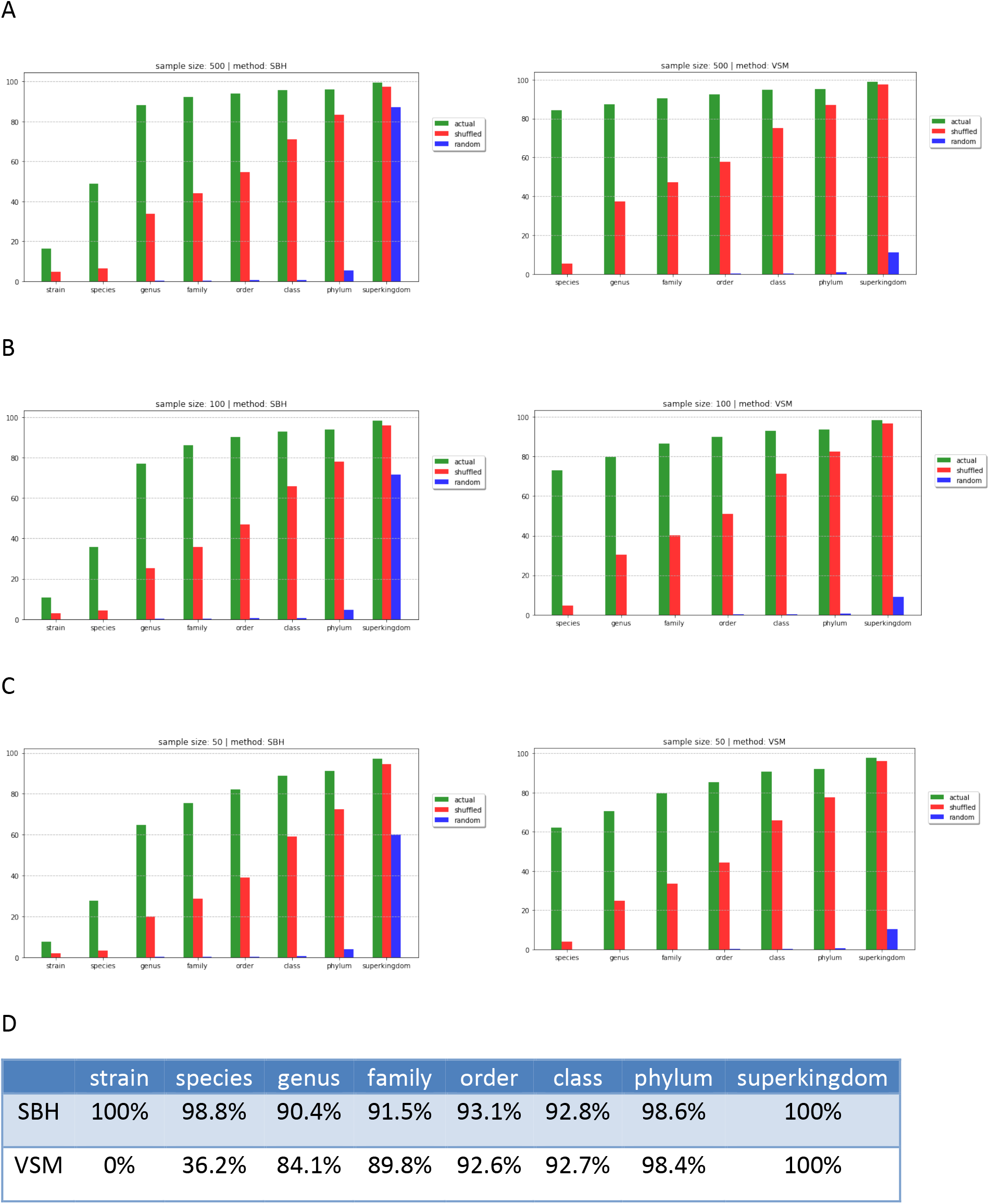
NCBI taxonomy benchmarking results for the following sample sizes: 500, 100 and 50 random gene translations, using SBH (left) and VSM (right). To obtain controls for each actual protein sequence used, two additional sets of decoy sequences mirroring the actual ones were created. These were designated as “shuffled” and “random”. Shuffled decoys were created by simple protein sequence shuffling creating a random order of initial amino acids i.e. the single amino acid frequency remained the same between actual and shuffled sequence, while the random decoy sequences used the same set of amino acids found in the actual counterpart and are of the same length, but the single amino acid frequency approached the normal distribution due to random sampling. (**A**) Randomly sampled protein sequences ranging from (**A**) 500, (**B**) 100 and (**C**) 50 proteins per taxon were decomposed into constituent 3-peptides, folded-in and used to construct queries to search against 54,526 taxa vectors in a species LSA model, built from the remaining sequences. (**D**) Table displaying percentage of taxa included in benchmarking tests for each taxonomic rank for both the SBH and VSM methods. Not all taxa have all ranks listed in the lineages and, since the VSM method includes the majority vote amongst the 5 best hits, it requires at least three taxa representatives for each query and for each rank. This excluded strain level from VSM benchmarking and significantly restricted the number of query candidates at species level being benchmarked.

These results suggested that random protein subsets of different sizes produced similar taxonomy mappings, which was surprising because of different set sizes and random selection of proteins in each set. To test this markedly different approach in comparison to sequence alignment, the LSA species model was compared to the widely used alignment-based method – BLAST [6]. Two non-overlapping, same-sized sets of sequences from 1,000 arbitrarily selected species were split 50/50 into training and test set (Supplementary Table 2). Taxonomy assignments were made the same as before for LSA. LSA utilizes multiple sequences forming a single query based on cumulative 3-peptide frequency opposed to BLAST, which performs solely sequence-to-sequence alignment, a concession had to be made where *a posteriori* only one good homology match between subject and query datasets was to be sufficient for taxonomic assignment made by BLAST (see Materials and Methods). Quite surprisingly, the alignment-free natural language processing method clearly outperformed BLAST (Fig. 2, A and B), although the comparison allowed for BLAST to take full advantage of coincidental homology between the query and subject protein sets.

**Fig.2.**
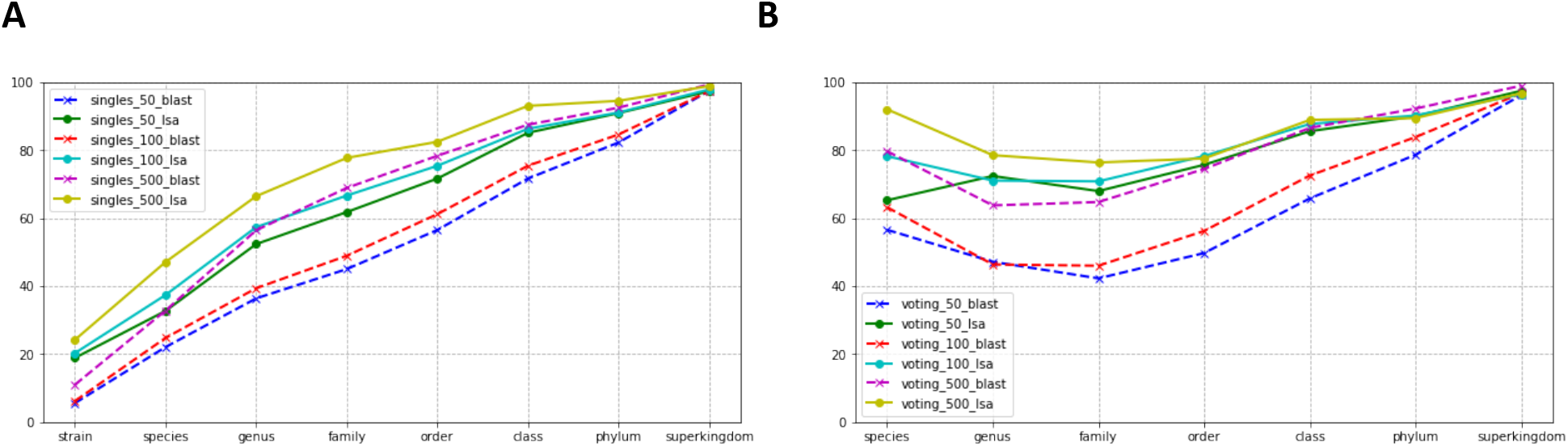
Taxonomy benchmarking results obtained by both LSA and BLAST for each of the three sample sizes (500, 100 and 50 randomly selected proteins) for both (A) SBH and (B) VSM methods. The X-axis differs between the two, because the VSM method was applied only to those taxa that had more than 3 representative organisms, therefore the lowest level (“strain”) is missing from the VSM results. Full lines represent LSA results and dotted lines represent BLAST results. Unlike results from the initial LSA benchmarking test, the VSM method gave a slight drop in correlation between species and family taxonomic levels, most likely because there were fewer organisms with 3 or more species or genus representatives in the 1,000 organisms selected (Supplementary Table 3)

This comparison of an alignment-based method with natural language processing, suggested that LSA was able to utilize some additional information content stored within heterologous protein samples that was not related to alignment-based homology.

### Even taxonomically restricted proteins exhibit semantic properties shared with their proteome

In order to further explore this possibility, coincidental homology between protein sets being compared had to be removed. To achieve this, taxonomically restricted sequences were used. Amongst these, orphan sequences are the most limited, restricted to only one or just a few species. These sequences are also interesting because they are frequently associated with novel phenotypes [13–15]. Therefore, the capabilities of LSA to extract knowledge from sequences beyond predictions obtainable using distant homology alignments [1] could be further explored. Declaring a gene or protein sequence as an orphan is very challenging because genomic databases are incomplete [16]. This is why the LSA approach was tested using sequences obtained by both stringent and relaxed scenario to account for the level of homology remaining within the sequence sets. The relaxed scenario included the NCBI Protein Clusters [17] dataset. All sequences within the proteomes of taxa used in the previous benchmarking experiment and listed in NCBI Clusters were regarded as orphan candidates and became part of the relaxed orphan dataset (Supplementary Table 4). This reversal in the way in which the Clusters dataset was created is analogous to alignment-based methods and E-value threshold criteria used for orphan sequence identification [18,19], and significantly reduces the overall homology between Clusters and orphan sequence sets used for the LSA model and subsequent queries. Because orphan candidates were those sequences not listed in the NCBI Clusters, this approach was designated as “relaxed”. In total, 3,913 unique species were present in both the NCBI “nr” based taxa proteome collection and NCBI Clusters, with > 100 orphan candidates. Exactly 100 proteins randomly selected from both Clusters and the orphan dataset represented each species twice, once as part of homology-rich Clusters and once as homology-poor orphans. Taxonomy-benchmarking tests comparing BLAST to LSA were performed in the same manner as previously described, with an additional step in which datasets were reversed. Despite favoring BLAST, the LSA predictions provided significantly better taxonomy benchmarking results (Fig. 3 A and B).

**Fig.3.**
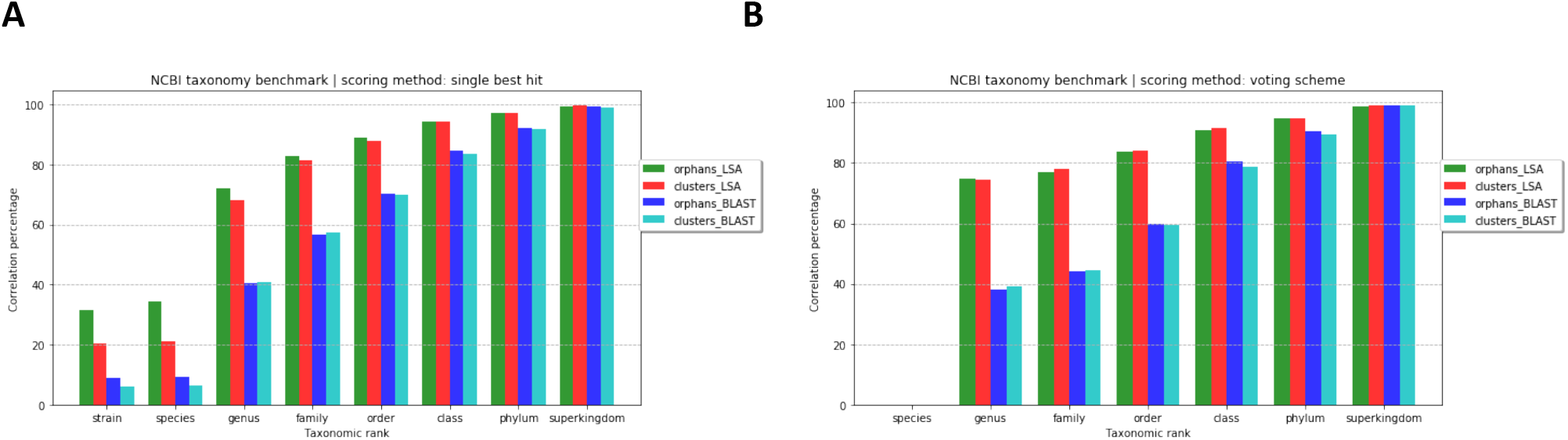
Relaxed orphan taxonomy benchmarking using (A) SBH and (B) VSM methods. The results were obtained in a reciprocal manner, in which the subject and query sequence sets were reversed. The combination in which the relaxed orphan dataset was used to build both the LSA subject vectors and blastp database, and where the Clusters dataset was used for query is denoted with “orphans_” prefix. The reciprocal combination in which the Clusters dataset was used to the build both the LSA model and blastp database, which were queried using the orphan dataset is denoted with “clusters_” prefix. Green and red bars display the correlation between the LSA benchmarking results and NCBI taxonomy database, whilst blue and cyan bars show the correlation between the results obtained using BLAST.

### Semantic properties of species proteins are completely independent of alignment-based homology

To completely rule out the effect of coincidental homology, all alignment-detectable homology was removed from the query sequence sets. To achieve this, a sample of 100 species was randomly selected, with all four kingdoms of life having at least one representative. In this sample, a two-stage filtering process removed all alignment-detectable homology (see Materials and Methods). In first stage, protein family members could be detected and removed using Hidden Markov models [20] and Pfam [21]. In the second step, extensive BLAST of the remaining sequences against the 54,526 species proteomes was performed to remove the remaining homology. What proteins remained after this filtering became part of the stringent orphan dataset (<u>Supplementary Dataset 1</u>). An LSA model was built using 54,526 species proteomes, with the orphan sequences removed. The benchmarking test was performed without BLAST comparison, because the second step of stringent filtering made this redundant. The results showed that 84% of species represented solely by orphan sequences, when used as a queries against LSA model displayed similarity to other related taxa (Fig. 4, A - E).

**Fig.4.**
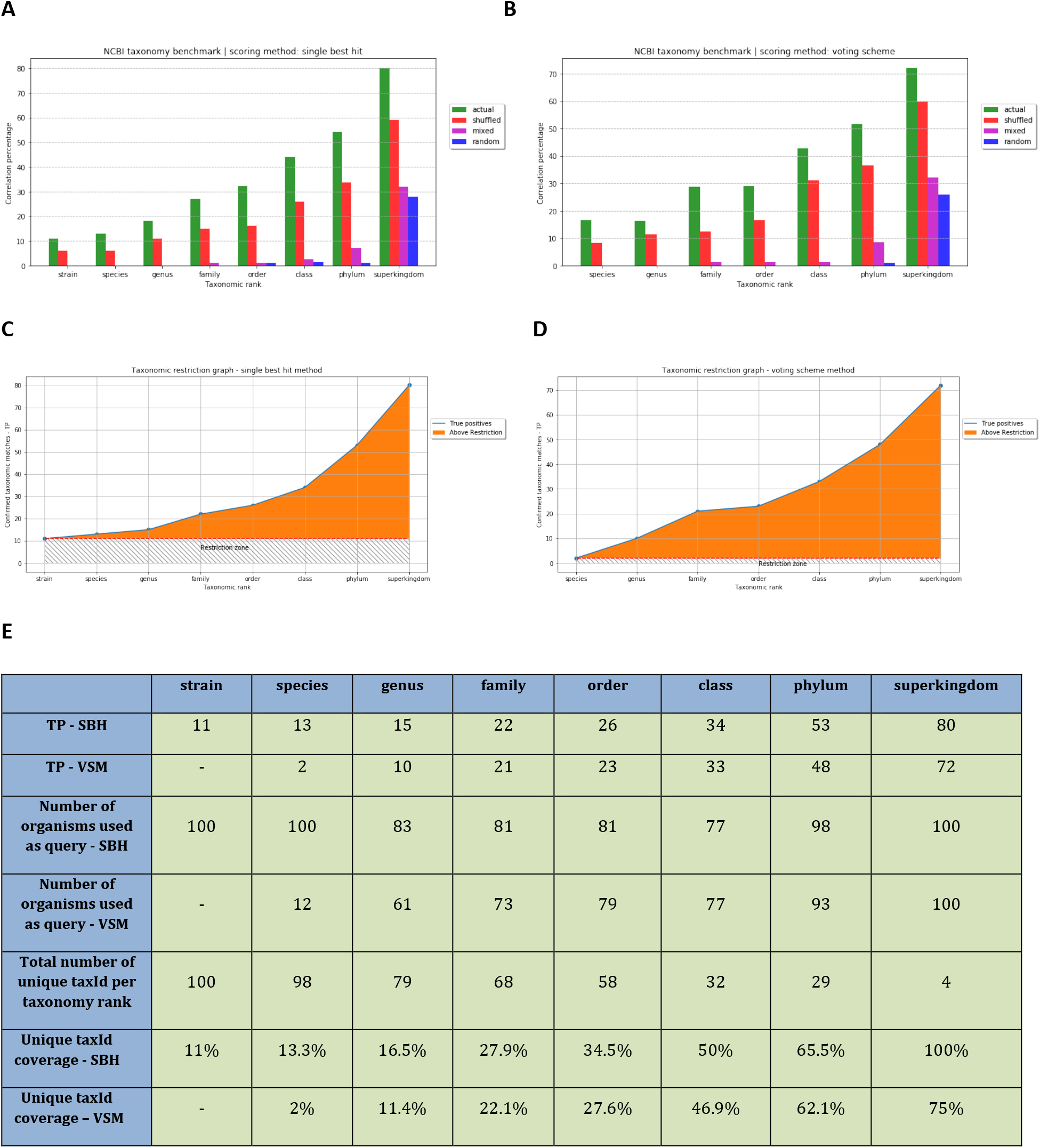
Stringent orphan taxonomy benchmarking results based on randomly sampled 100 organisms. Results for (**A**) SBH and (**B**) VSM methods. For each orphan sequence, three decoy sequences were included in the analysis. These decoys were used to create “shuffled”, “random” and “mixed” datasets. Shuffling the entire orphan dataset and then re-populating each subject taxon with the same number of proteins taken from this collective mix made the “mixed” dataset. The total number of positive taxonomic matches (TP) for each rank is shown for (**C**) SBH and (**D**) VSM methods. Horizontal dashed lines at the bottom of each plot denote the expected taxonomic recognition threshold (as defined by the query sequence being restricted on the lowest taxonomic level) (**E**) Tabular overview of results where the first two rows display TP count at each taxonomic level. The third and the fourth rows show the number of organisms available as a query at each taxonomy level. The fifth row gives the number of unique taxonomic identifiers amongst the 100 selected organisms – representing the taxonomic diversity of the sample. The sixth and seventh rows display the percentage of taxonomic diversity within query organisms that was successfully covered by positive matches (total unique taxId per rank / unique taxId per positive matches) for both SBH and VSM methods.

To rule out the possibility of obtaining comparable results by chance, “mixed” and “random” decoy sequences were used as controls (Fig. 4, A and B). The majority of positive taxonomic assignments were achieved beyond the homology restriction level (Fig.4, C and D). This strongly implied that LSA established relationships between orphan proteins and related species proteomes, based on taxonomy and in the total absence of alignment-detectable homology. This common taxonomy-related denominator was further highlighted by the “mixed” query set, which was constructed from sequences of mixed origin. Unlike “shuffled” or “random” decoy sequences, by using actual sequences of mixed taxonomic origin, real information content of the protein sequences was used but the taxonomic contribution was removed. Mixed sequence results were most comparable to those of random decoys and performed significantly worse than actual or shuffled sequences. This confirmed the importance of 3-peptide frequency patterns within species proteomes and indicated that all proteins sharing a proteome context include additional information content, even in the complete absence of alignment-based homology. True orphans are considered to be a subset of taxonomically restricted proteins, which are unique to a specific taxonomic level. In contrast to non-orphan proteins, orphans are believed to be unique to a species, however these results (Fig. 4 C and D) indicate that this is not the case when alignment-free similarity methods are used.

### More than just bias – species proteomes exhibit distinct 3-peptide frequency signatures

Different species have different codon usage preferences. A major difference between codon frequency and amino acids frequency is that different codons can code for the same amino acid, while different combinations of amino acids result in different molecules. Codon usage bias is a well-accepted phenomenon, but the consequence of biased DNA translated to biased proteins on a genome wide scale is somewhat less well known [22]. Different species may have different preferences for specific amino acids codons and consequently, these codons will occur at uneven frequencies in the population causing preferences for specific amino acids to differ [23,24]. Following this rationale, LSA was used to investigate the possible existence of more systematic biases between entire collections of species proteins - proteomes. This was tested using both random sequence sets sharing solely coincidental homology and orthologous sets sharing maximal homology across species. Shuffling sequences is commonly used to create decoy datasets when estimating false discovery rates [25,26]. Since LSA can utilize multiple sequences as query and notionally “amplify” signals coming from individual sequences, this method might not suffice. To test whether this was the case, each 3-peptide in the “actual”, “shuffled” and “random” datasets was counted for > 27 million proteins, from each of the 54,526 taxa used throughout this study (Fig.5 A).

**Fig.5.**
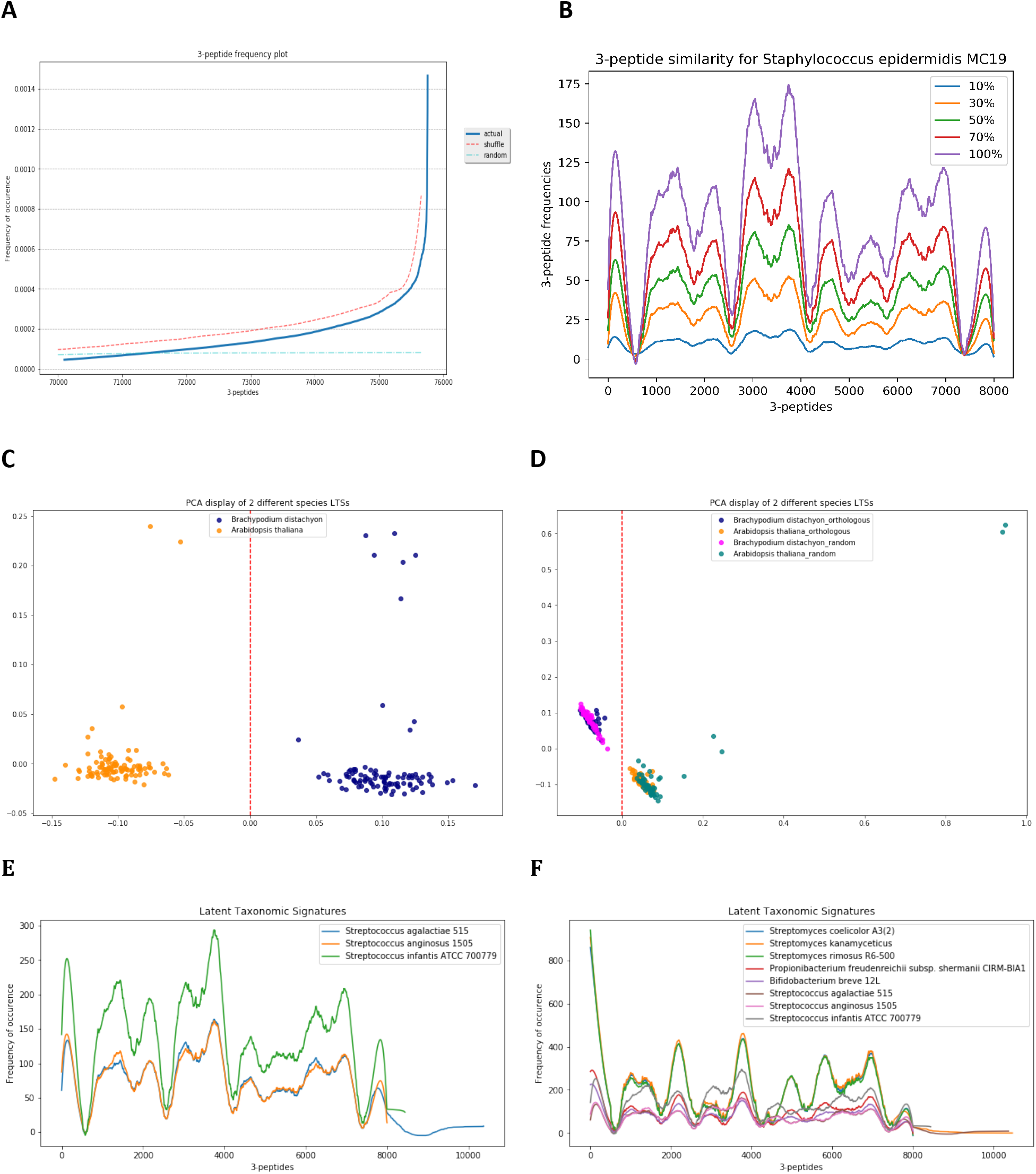
All of species proteins exhibit taxonomy-related signatures. 3-peptide frequency distribution and selected taxa 3-peptide frequency plots. (**A**) 3-peptide frequencies were calculated from a sample of 27,263,000,00 proteins. The frequencies (Y-axis) were calculated for “actual” (blue), “shuffled” (red) and “random” (cyan) sequences. The X-axis represents 3-peptides sorted by decreasing frequency using the “actual” sequence dataset for reference. Shuffled and random 3-peptide frequencies were shifted equally on the x and y-axis to be discernible. Only the most frequent 5,658 3-peptides are being displayed. (**B**) Random protein samples containing a different percentage of a complete proteome displaying a “signature” in the form of a fractal pattern, as seen here in a randomly selected bacterium. (**C**) Two random plant species proteomes each represented by a random selection of 100 proteins by 100 iterations. Each iteration was folded into LSA vector space and displayed using a 2-dimensional PCA plot (**D**) The same two plant species, each represented 50 times using random 100 proteins (with virtually no homology between two species proteins sets) and another 50 times by a different set of 100 shared orthologous proteins (with the highest possible degree of homology between two species protein sets). (**E**) 3-peptide frequency plots displaying similar signatures of 3 species of Streptococcus. (**F**) 8 different taxa “signatures”, 3 Streptococcus species, 3 Streptomyces species, 1 species of Bifidobacterium and 1 species of Propionibacterium. A Savitzky-Golay filter was used for smoothing of all plots displayed. Closely related species have more similar signatures.

When displayed on the same plot, the 3-peptide frequency of “shuffled” sequences exhibited a similar pattern to the “actual” sequences, strongly implying that shuffling was not sufficient for the collection of proteins to lose the taxonomic signature. This was confirmed using a two-sample Kolmogorov-Smirnov test (KS statistic=0.02474 and p_value=0.062596 for α=0.05 with critical value D_crit=0.0255695) calculated on 5,658 of the most frequent 3-peptides from the “actual” protein dataset. While sequence shuffling can make a single sequence sufficiently random for alignment-based methods, it was not sufficient for LSA using more than one sequence to construct meaningful queries, even in the absence of alignment. This explained the difference between the “shuffled” decoy results when compared to results for the “random” and “mixed” datasets. The 3-peptide frequency pattern was an integral feature exhibited by all proteins making up a proteome (Fig. 5B). Whilst the proceeding benchmarking results (Fig. 1 A, B and C) indicated these signatures were species specific and robust, this had to be tested by repeated subsampling of different proteome samples. Although each sample was a completely different mixture of species proteins, the boundary between species remained stable (Fig. 5C). Perhaps the most striking confirmation of robustness was obtained when orthologous sequence samples were mixed with random sequences (Fig. 5D). Even when same sets of orthologs were used, the species vector clustering remained stable despite ortholog protein family relatedness. Smoothed histograms of 3-peptide frequency data were used to demonstrate signature similarities between closely related taxa (Fig. 5 E and F). These proteome-distributed features were designated ‘Latent Taxonomic Signatures’ (LTSs) because is the information is both hidden and discernible from information obtained by alignment-based homology. The discovery of LTSs posed an intriguing question, how did genes belonging to different gene families manage to produce a feature distributed amongst different proteins that constituted species proteomes?

### Are taxonomic signatures resulting from context dependent evolution?

Besides originating from related organisms, it is difficult to explain any commonality between proteins displaying LTSs. The same LTSs were observed in heterogeneous protein subsamples encoded by genes from different gene families that shared only genome and proteome context, but no alignment-based homology. This suggested that all protein-coding genes within taxa evolve in a concerted manner, dependently of taxonomic context. Previous results demonstrated that not even orphan sequences were exempt from this (Fig. 4). However, these datasets represented heterogeneous collections of functionally and structurally unrelated species proteins. Although random sampling is usually the method of choice for testing hypotheses, to further validate universality for the proposed constraint, the occurrence of LTSs was also established in non-random, homogenous protein samples. To accomplish this, protein families were used to collect sequences and further sub-divided into taxonomic groups, with each group folded-into a single LSA vector to represent the group. Each of these vectors was created using 3-peptides from highly homologous sets of proteins characterized by significant sequence similarity and a well-defined alignment. These vectors represented both protein families and contributing taxa, and were compared pairwise within the same protein family e.g. “intra-class”, and to the entire collection of species vectors, based on whole proteome 3-peptide frequencies e.g. “inter-class”. Cosine based comparison was performed with two possible outcomes, “selfish” or “altruistic”. A null hypothesis assumed that there was no relationship between the taxonomic origin of the proteins and the outcomes. Hence, the 3-peptide frequency pattern for any given protein family taxonomic group was expected to be more similar when compared “intra-class” than “inter-class”. A high degree of homology within protein family versus no-homology within a collection of proteins contributing to a proteome would certainly have indicated this. To test this hypothesis, two control datasets were added, the “mixed” control dataset, with sequences randomly sampled from a protein family (mixed taxonomic origin, but possibly biased) and, being completely taxonomically independent, an “HMM” control dataset where protein sequences were replaced by the consensus sequences generated from a protein family HMM model [20],. In cases where the observed intra-class similarity was greater than any of the inter-class similarities, the outcome was regarded as “selfish”. In the opposite scenario, when one or both inter-class similarities prevailed, the outcome was regarded as “altruistic”. The results of these comparisons indicated more similar and abundant inter-class outcomes than intra-class similarity outcomes (Fig. 6A).

**Fig.6.**
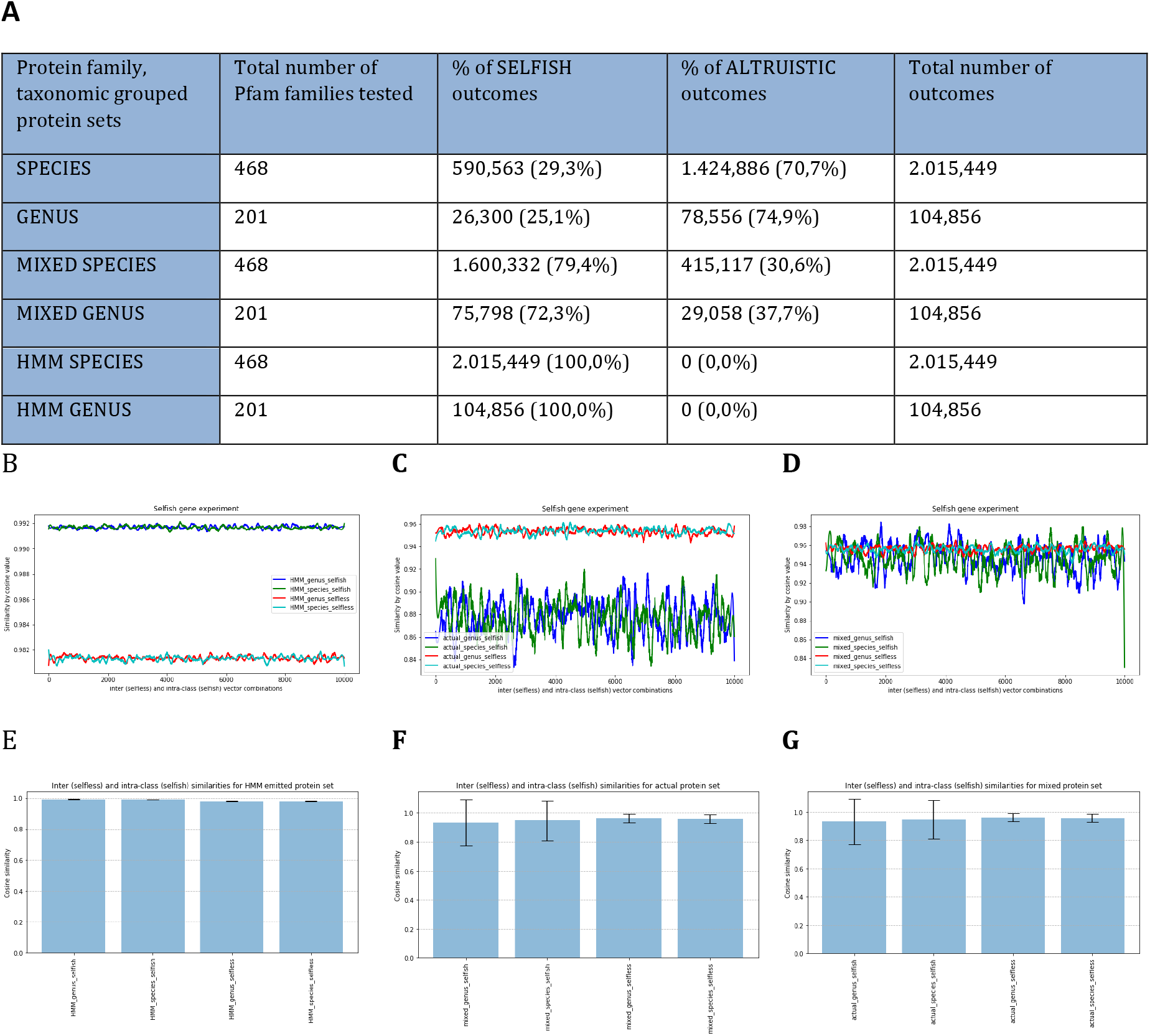
Taxonomy context dependence tested on 468 Pfam protein families in which proteins were grouped according to species, and 201 families where proteins were grouped according to genus (Supplementary Table 5). (A) Table summarizing the experimental outcomes for “actual” (no prefix), “mixed” (prefix MIXED) and HMM (prefix HMM) datasets. HMM consensus sequences produced exclusively “selfish” outcomes, whilst the majority of actual sequence outcomes were “altruistic”. (B) HMM taxonomy independent inter- and intra-class cosine similarity plots constructed from a random selection of 10,000 family-taxa combinations showing best recorded intra-class (selfish) and inter-class (altruistic) cosine values for each of the 10,000 pairwise comparisons. Selfish outcomes are characterized by significantly higher recorded cosine values as expected. (C) Cosine plot from a random selection of the 10,000 pairwise family-taxa actual protein set combinations representing “selfish” and “altruistic” outcomes. Selfish outcomes are characterized by significantly lower and more variable recorded cosine similarities compared to “altruistic” outcomes, which indicated an evolutionary constraint imposed by species. (D) “Mixed” species inter- and intra-class cosine plot for a random selection of 10,000 pairwise comparisons. (**E**) HMM consensus sequences error bar plot. (**F**) Mixed proteins error bar plot. (**G**) Actual proteins error bar plot. In all error bar plots, the height of the bars is the mean intra-class and inter-class similarity represented as a cosine value, for both species and genus grouped sequences. The error bars are a +1/-1 standard deviation about the mean. For the inter-class similarities, an average of two best cosine values obtained by comparison with species LTSs are given. For all three datasets (HMM, actual and mixed), all possible pairwise combinations were used in the table and only 10,000 random combinations for the plots.

Intra- and inter-class cosine similarity measures for all three datasets gave the same frequency distributions based on Mann-Whitney U Test (p = 0). A Chi-Squared Test on the outcomes from all three datasets (Fig.6, A) rejected the null hypothesis (p=0 with a significance level of 0.05). Experimental outcomes obtained from the HMM dataset represented a gene-centered view expectation. However, in the case of actual sequences, this was not the case. Inter-class similarity outcompeted intra-class homology (Fig.6 B and C). This was further supported by mean values from the cosine similarity measures. The mean cosine value for the HMM control dataset was significantly higher (0.992) for both species and genus groups in selfish outcomes, compared to 0.981 for the altruistic outcomes. Conversely, actual sequences afforded significantly lower mean cosine values, with 0.879 (species) and 0.886 (genus) for “selfish” outcomes, compared to “altruistic” outcomes of 0.954 (species) and 0.960 (genus). The mixed dataset was more aligned with the HMM dataset outcomes. However, because some families had taxonomic biases, the outcomes were also largely mixed (Fig.6, A and D). Data used for this experiment were evaluated by constructing error bar charts to display inter- and intra-class cosine similarities for all three datasets (Fig. 6, E, F and G). These charts suggested that the amount of uncertainty was highest in the actual dataset, when “selfish” outcomes were allocated. When “altruistic” outcomes were observed, the degree of uncertainty was much lower and comparable with the control HMM dataset.

## Discussion

Statistical analysis of homology between biological sequences is a standard approach to identify conserved sites and motifs that correlate with biological knowledge, such as structure and how this relates to function [27]. Function related knowledge is commonly assigned solely based on sequence alignments. However, this kind of homology based annotation provides limited insight, for example, complementation studies between mutations in a protein can infer interactions between amino acid residues not readily observed within an alignment [28]. Thus, a longstanding problem in bioinformatics is inferring the information encoded in a biological sequence, when there is little or no sequence homology. A promising new field of research in biology is natural language processing, with the original task of determining the meaning of a word (i.e. semantics) from the contexts in which the words appear [3] being translated to biological sequences. Natural language processing and distributional semantics have increasingly been adopted in biology, as an alternative to alignment-based homology [29–31]. In this study, Latent Semantic Analysis (LSA) was applied to a large number of different species proteomes, spanning complete range of taxonomic diversity found in current databases. This LSA based species model identified conceptual content from heterogeneous collections of proteins contributing to proteomes, by establishing associations between 3-peptides that occurred in similar contexts. The resultant 3-peptide signatures were discernible features characterizing each species proteome, where protein subsets of a proteome produced signatures that were small replicas of the whole species proteome ones, resembling self-similar property of fractals. A major difference between this approach and sequence alignment is that an alignment is primarily an analytical method, restricted to comparisons between contiguous sequences and subsets of conserved residues. Conversely, natural language processing and LSA represent a more holistic approach, which places dispersed motifs into the context of entirety. Consequently, species can be modeled as a collection of random sequences, as opposed to being reduced it to a single gene or protein family representative. The alignment is not a prerequisite for this kind of modeling, since vectors that represented species proteomes provided valid taxonomic knowledge, even in the total absence of meaningful sequence alignment (Fig 3. A and B, Fig 4. A and B). Evolution is biased towards selecting conserved amino acids that are consistent with fitness [32]. Taking a gene-centered view, evolutionary success is directly linked with the ability to generate as many near-identical copies in a population as possible. Hamilton mathematically set foundations for this reproductive fitness as a measure of evolutionary success in his “Inclusive Fitness” theory of 1964 [33]. Results presented herein suggested that notion of a successful replicator encoding solely self-contained information was incomplete. No gene or protein exists as an atomic entity but resides within a taxonomic context. In such a context, transgenerational transmission of a gene or protein would unlikely be successful without additional information that describes reciprocal interactions between other genes comprising the genome and equally, other proteins representing the proteome. Thus, the total information content in a genome and proteome is inextricably linked by integration between these salient features. LTSs indicated a bi-directional relationship between the species and protein content within the species. In this relationship, genes affected species but the genomic and proteomic context of species constrained individual gene and protein evolution. A concerted change would be required in order to preserve both primary function and the majority of contextual information. Restrictions on the independence of chromosome evolution and examples of concerted evolution phenomena are well known [34,35], but have been less well demonstrated at the level of entire genomes or proteomes [36]. Another concerted, but only recently explained process, is conservation of replication timing order, which has been shown to orchestrate the global epigenetic state of individual cells. The principle behind this process is the continuation of previously established functional interpretation of the information stored in the genes [37], which follows the rationale described in this study.

Beside random protein samples, even highly homologous groups of proteins from different protein families displayed greater affinity towards a proteome context than towards constituent family sequences, when modeled semantically. Finally, the most surprising confirmation of this constraint came from orphan protein sequences. Based solely on sequence alignment, orphan sequences are considered both taxonomically restricted and *de novo,* with no apparent relatedness to established protein families. Results presented in this study indicated that even the most stringently defined orphan sequences were not taxonomically restricted, since they were also found to posses LTSs. This placed orphan sequences in proximity to taxonomically related organisms outside the alleged “restriction zone” (Fig 4. C and D). Since protein evolution is a direct consequence of changes and selection of DNA polymorphisms and mutations, the real question is what could have caused both constrained and concerted evolutionary pattern observed by LTSs? Thus far, the only plausible evolutionary process that acts to maximize genomic and phenotypic cohesiveness linked to speciation is Molecular drive. Understanding how information in a genome and proteome are inextricably linked to a context, and how context is conserved between subsequent generations warrants further experimentation.

## MATERIALS AND METHODS

### Obtaining data

All protein sequences were collected from NCBI “nr” database for each of the species included in LSA model (https://www.ncbi.nlm.nih.gov/refseq/about/nonredundantproteins/). Proteins were linked to taxa using python programs we have written, which relied on accession to taxId mapping files provided by NCBI Taxonomy database (https://ftp.ncbi.nlm.nih.gov/pub/taxonomy). NCBI Taxonomy database was used as a reference taxonomy database to benchmark LSA against. All used protein data was stored in a local copy of a simple relational database we made using SQLite and python scripts. A Python framework for the analysis and visualization of trees (http://etetoolkit.org/) was used for dealing with the NCBI Taxonomy database [38]. It was used to make a local copy of the NCBI Taxonomy database, and it was later used in our programs to convert from taxId to scientific names (and vice versa) and to establish taxonomic ranks, names and lineages. For relaxed orphan dataset, NCBI Protein Clusters repository was used (https://ftp.ncbi.nih.gov/genomes/CLUSTERS/) in combination with the local copy of “nr” based SQLite relational database. For stringent orphan dataset, we have downloaded current release of Pfam-A HMM models (ftp://ftp.ebi.ac.uk/pub/databases/Pfam/current_release/Pfam-A.hmm.gz) and used HMMER v.3 (http://hmmer.org/) to scan protein family HMM models against previously selected proteome sequence sets which were extracted from the local database and written in form of simple multi-FASTA protein files. Pfam-A dataset was also used for the “selfish” vs “altruistic” genes experiment as a source of protein family HMM models.

### Transforming proteomes into “words”

Let *AA* be set of 20 natural amino acids. Let *p* = *a*_1_*a*_2_*a*_3_ … *a*_*n*_represent protein, where *a*_*i*_ ∈ AA and *n* is length of protein.

We define triplet *t*_*i*_ as

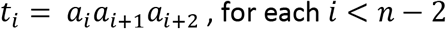

### Truncated singular value decomposition

After *m*×*n* matrix *A* has been created and properly weighted, a rank-k approximation (*k* ≪ min (*m*, *n*)) of matrix *A*, *A*_*i*_, is computed using truncated singular value decomposition (T-SVD). Singular value decomposition is a technique closely related to eigenvector decomposition and factor analysis, used to model the associative relationships. This allows for both integrative proteome comparison and topic modeling of 3-peptide motifs within protein sequences. With truncated SVD, matrix *A* is factored into the product of 3 matrices: *m*×*k* term-concept vector matrix *U*_*i*_, a *k*×*k* singular values matrix ∑_*k*_ and *n*×*k* document-concept vector matrix *V*_*k*_. Singular value decomposition allows the arrangement of the space to reflect the major associative patterns in the data, and ignore the smaller, less important influences. As a result, terms (in our case 3-peptides) that did not actually appear in a document may still end up close to the document, if that is consistent with the major patterns of association in the data. The truncated SVD captures most of the important underlying structure in the interrelation of terms (3-peptides) and documents (proteomes) and at the same time much of the noise that causes poor retrieval performance is eliminated. In the reduced space, semantically related terms and documents probably lay near each other since the SVD attempts to obtain the fundamental, semantic structure of the term-document (3-peptide – proteome) space. Matrices and process are illustrated below (Fig. 7).

**Fig.7.**
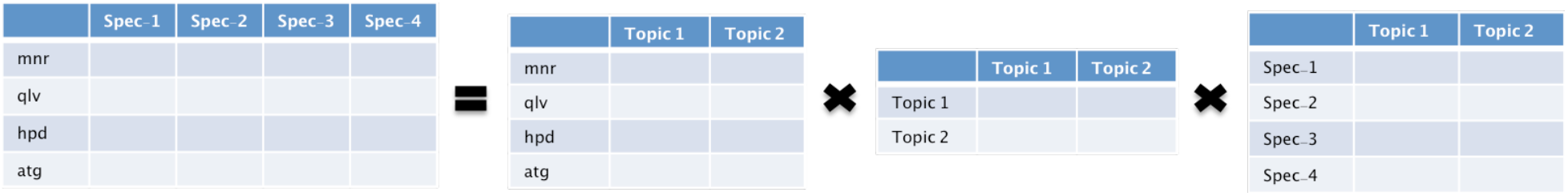
An overview of T-SVD of 3-peptide (terms) – species proteomes (documents) matrix. With truncated SVD, 3-peptide - species matrix is factored into the product of 3 matrices in the following order (left to right): m×k 3-peptide-topic (e.g. concept) vector matrix U_i_ followed by k×k singular values matrix ∑_k_ and final n×k species-concept vector matrix V_i_ which is the source of our species proteome vectors.

### Latent semantic analysis

Typically, database is organized so that information is retrieved by literally matching exact terms in documents with those of a query. Since there is usually great amount of common 3-peptides in different species proteome, the literal terms in a query may not match those of a relevant document. A better approach would allow users to retrieve information based on the meaning of a document. Latent Semantic Analysis (LSA) [2] is an extension of the vector space retrieval method in which the dependencies between terms are explicitly considered in the representation and exploited in retrieval. This is done by simultaneously modeling all the interrelationships among terms and documents. The LSA is a retrieval method that builds upon the prior research in information retrieval - Vector Space Model, using the Singular Value Decomposition (SVD) [39] to reduce the dimensions of the term-document space. In vector-space retrieval, a document is represented as a vector in *m*-dimensional space, where *m* is the number of terms in the lexicon being used. Each component of the vector reflects a concept associated with the given document. More formally, let *m* be the number of words (3-peptides) in the dictionary, *n* the number of documents (species) and *d*_*i*_ ∈ ℝ^*m*^, *i* = 1, …, *n*, tf-idf representation of *i-th* document. Then the term-document matrix *A* has the following form:

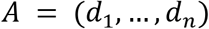

*A* is usually very large and very sparse matrix since the number of terms in each document is significantly less than the number of terms in the entire document collection. Once a term-document matrix is constructed, local and global weightings are applied to increase or decrease the relative importance of terms within documents. We must also represent the query by the tf-idf scheme, the vector *q* ∈ ℝ^*m*^. We would now like to calculate the singular decomposition of the matrix *A* = *U*∑*V^T^* which has the following shape:

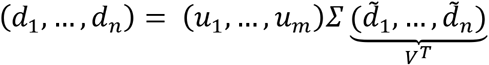

Suppose we construct the matrices *U*_*k*_, ∑_*k*_ *i V*_*k*_. The representation of the document (species) *d*_*i*_ in the vector space {*u*_*1*_, …, *u*_*k*_} is

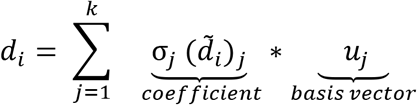

The singular value of σ_*i*_ represents the importance of *i-th* basis vector, *u*_*i*_. The columns of the matrix ∑_*k*_*V*_*k*_^*T*^ T represent the coefficients of documents (species) in the newly created lower dimension space, i.e. latent space. In order to be able to compare the query (a set of proteins sharing a proteome) with documents (species), we must represent the query within the latent space. Namely, we look for the coordinates of this vector in the space spanned by the columns of the matrix *U*_*k*_. If we denote by 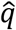 projection of the vector q on that space, then we know that:

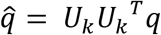

Thus, the vector *U*_*k*_^*T*^*q* represents the coefficients of the vector q in the latent space, i.e., the space spanned by the columns of the matrix *U*_*k*_. The coefficients of other documents (species) in the latent space are given by the columns of the matrix ∑_*k*_*V*_*k*_^*T*^. In order to avoid explicit computation of the matrix ∑_*k*_*V*_*k*_^*T*^we define

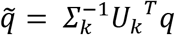

This is also called *folding-in* method [40]. Once the query is projected into the term-document space, cosine similarity measures is applied to compare the position of the new-created query to the positions of the terms or documents in the reduced term-document space. We compare the query *q* with the documents (species) by comparing 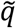 with the columns of the matrix *V*_*k*_^*T*^. In the end, we count

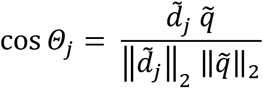

for each *j* = 1, …, *n*.

### Practical application of natural language processing

All LSA related modeling was performed using Gensim. Gensim is an open-source library for unsupervised topic modeling and natural language processing, using modern statistical machine learning. Gensim is implemented in Python and Cython (https://radimrehurek.com/gensim/) and was installed as part of a python virtual environment with all of the dependencies. Beside Gensim, numpy (https://numpy.org/) and scipy (https://www.scipy.org/) were used for vector operations and statistical analyses. Figures were created using matplotlib (https://matplotlib.org/) and Jupyter Notebook, which were in the same virtual environment Gensim was installed in. All of the libraries and code written was deployed on x86_64 Debian based Linux Ubuntu 18.04.5 LTS local server machine with 2 Intel Xeon E5-2650 2.00GHz CPUs, 8 cores per socket and 2 Threads per core and 264 GB RAM. This server also hosts the web accessible version of LSA based species model. This model was deployed as a web application making it easy for anyone to explore taxonomic context of selected proteins, which is simple to use, requiring no programming or NLP knowledge and no alignment-based homology in the proteins being compared: http://matrix.pbf.hr/.

### LSA model of species

LSA uses a term-document matrix, which describes the occurrences of terms in documents. In our model, we replace documents with taxa, and words (also called terms) with all occurring 3-peptides extracted from proteins within species proteomes. In order to transform proteomes into LSA vector space, we decomposed the entire constituent proteins into 3 consecutive amino acid residues e.g. 3-peptides. This is the same length BLAST [6] uses as default “word” size for a protein sequence. In our case, this was accomplished by first lowercasing all the protein sequences followed by decomposition into consecutive 3-peptides using a sliding window method, with frame equaling 3 amino acids and “sliding” step equaling to one (Supplementary Figure 1A). Applying dimensionality reduction (SVD) we have embedded distributional information on all occurring 3-peptides into 400-dimensional LSA vector space. This allowed us to compare entire species proteomes and explore their distributional semantic similarity by correlating it with established taxonomic classification using cosine similarity as metric (Supplementary Figure 1B). Final result aggregates proteome information on tens of thousands of taxa collected from both NCBI Taxonomy database and non-redundant protein database. There are more taxa in these databases; however, we have included only those represented with more than 100 proteins. In order to handle big protein data, we restricted the maximal number of protein sequences to be included in the LSA model to no more than 5,000 proteins per taxon. This means that for all taxa which have proteomes represented with more than 5,000 proteins, 5,000 proteins were randomly selected, while for the rest of the taxa having protein datasets between 100 and 5,000 proteins, entire protein sets were tokenized (Supplementary Table 1). This model is an example of the “bag of words” model [3], which disregards 3-peptide ordering but keeps their multiplicity. On this model, first a term frequency - inverse document frequency (TF-IDF) weighing scheme was implemented [41]. During TF-IDF transformation, 3-peptides which were rare in the training corpus had their weights increased without changing the vector dimensionalities. Finally, we have used LSA to transform from TF-IDF weighted space into a latent space of a lower dimensionality. In our case, we have chosen 400 dimensions empirically [42]. The entire process is summarized below (Fig. 8).

**Fig.8.**
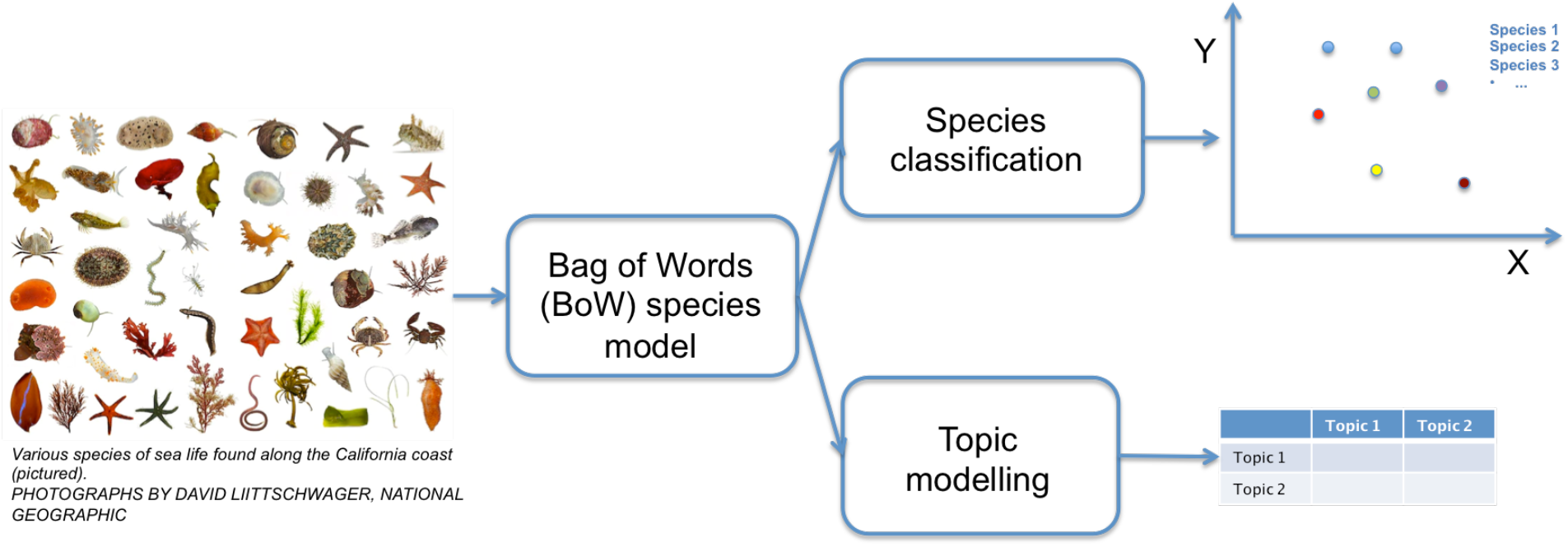
An overview of LSA based species model. Species proteomes are represented as “bags of 3-peptides”. First step of transforming species proteomes into 400-dimensional vector space (here depicted in simple 2D) includes building a dictionary, which simply counts each 3-peptide occurrence in each species proteome. In the second step, 3-peptide counts are weighted by TF-IDF that gives more weight to those 3-peptides occurring frequently within species proteome and seldom within the rest of the species proteomes in the initial matrix. Finally, SVD is used to transform weighted 3-peptide counts into 400-dimensional species vector space.

### Taxonomy mapping (benchmarking) tests

Random subsampling was used to create proteome subsets used for taxonomy mapping. In all four cases, the actual proteins being tokenized and folded into query vectors were searched against the LSA species model and two different methods have been applied in order to assign taxonomic information to query vectors. First method named “single best hit” (SBH), directly maps taxonomic information belonging to most similar subject vector (by cosine value) to the query. If this assignation correlates with the NCBI Taxonomy taxId of query proteome it is regarded as positive match, otherwise it is negative. Second method is the “voting scheme” (VSM), which is based on taxonomic information from 5 highest-ranking subject vectors (based on cosine) and used in a voting scenario. This scenario requires unambiguous taxonomic majority among the highest-ranking subjects. If this majority correlates with NCBI Taxonomy taxId for the query proteome it is regarded as positive match otherwise it is negative (Supplementary Figure 1C). In case there is no majority vote, the benchmarking result for this query is regarded as negative. Other difference between SBH and VSM methods is the total number of valid query proteomes. In case of SBH the query space is the same as the number of taxa in initial LSA matrix (subject space). In case of VSM, query size is somewhat smaller. The reason for this discrepancy is the fact that the voting scheme relies on majority vote. This majority in worst-case scenario means 3:2 ratio between subject taxa and therefore, query space is restricted only to those taxa, which have at least three representatives in all taxonomic ranks benchmarked. This also implies that VSM will not be including the lowest “strain” taxonomic level, into benchmarking results, unlike SBH. Since cosine value is used as taxa proximity metrics, we have created two new proximity values called “adjacency score” and “similarity interval”. These values serve the role of making taxonomic assignations based on cosine similarity between query and subject taxa vectors simpler and more intuitive. Adjacency score is calculated using formula below:

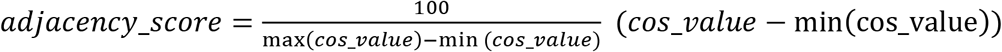

Because cosine range of [0,1] is very narrow, when querying LSA species model we get very dense distribution of subject cosines. This is why in our web application “Matrix of Life” (http://matrix.pbf.hr) only the most similar 100 subject vectors taxonomy information is being displayed. Therefore we decided to “stretch” obtained cosine values and present them on a scale ranging from 0 to 100. The formula above transforms cosines into adjacency scores in a way that the minimal reported cosine maps to 0 and the maximal one to 100. Similarity interval is even simpler to interpret. Essentially, we have divided the [min(cos_value), max(cos_value)] interval on 5 equal parts and counted subjects contained within each part. All subjects within this new interval get a score ranging from 1 to 5 (1 for being the first interval, 5 for being in the last).

### Comparison between LSA and BLAST

For BLAST comparison, we have used locally compiled and installed NCBI BLAST 2.9.0+ with “makeblastdb” application to produce BLAST databases from the same proteome sets, which were used to construct LSA vector spaces. Same query proteome sets were used for both LSA queries and “blastp” queries. In order to compare LSA benchmarking results using previously described SBH and VSM methods for taxonomic assignations we have decided to use BLAST favoring conditions. For comparison with SBH, in case of BLAST obtained results, best E-value (lowest) overall HSP (High-scoring Segment Pair) obtained by searching all query proteins against all subject taxa proteins was used. For comparison with VSM method same “blastp” search was performed, but in this case best five HSPs overall (based on E-values) were used to infer votes and assign taxId to query if there is a majority vote. This concession had to be made because unlike LSA, which utilizes information stored within multiple query sequences, performing a single search across all subject vectors, BLAST is limited to information stored within a single, contiguous sequence and performs M×N_sample_ pairwise comparisons for each query, N_sample_ being the sample size (500, 100 or 50 sequences) and M=N_taxa_×N_sample_, where in this case N_taxa_= 1,000. This concession meant that for BLAST only, the best overall pairwise-comparison was used to make taxonomic assignations, and all other matches were disregarding.

### Stringently defined taxonomically restricted (orphan) proteins

Using Python scripts we wrote, two-stage orphan candidate filtering was performed on randomly selected 100 taxa proteomes. Since the taxonomic composition of the initial dataset is dominantly Bacterial (92% of taxa), while the other three superkingdoms represent 5,5% (Eukaryota), 2,5% (Archaea) and 0,5% (Viruses). We have sampled 100 taxa more equally from all four superkingdoms to ensure they all have their representatives. First stage orphan candidate filtering included screening for protein family members using Pfam database. Pfam is a large collection of protein families, each represented by multiple sequence alignments and hidden Markov models (https://pfam.xfam.org/). For this screening, Pfam-A collection of HMM profiles was used since it is based on the same non-redundant (“nr”) protein sequences from the NCBI ftp site (https://ftp.ncbi.nlm.nih.gov/refseq/), which we used to create LSA species model. Screening was performed using HMMER “hmmscan” search program and Pfam-A collection of HMM models to search against each of the 100 taxa proteomes in 100 selected taxa. Default “hmmscan” E-value threshold of 10.0 was used. Results of this search were saved in tabular formatted files, which were parsed, and only proteins with no detectable protein family signal were retained (proteins which were not included in the “hmmscan” report). These proteins were selected as first stage or “candidate” orphans. In the second stage “candidate” orphan proteins from the first stage were used as queries in a “blastp” search. This second step served to remove all those proteins, which passed first stage, but still might have homologs that are not assigned to particular protein family. In order to construct blast database for this second step, from each of the 54,526 taxa having more than 1,000 proteins, 500 proteins were randomly sampled and written as a single FASTA file that was turned into a “blastp” searchable database using “makeblastdb” command. In this manner we could perform BLAST search with all the first stage candidate orphan proteins in reasonable time on our server machine, without compromising opportunity to catch stray homologs within entire taxa range we plan to query against. The most commonly used “blastp” E-value threshold for orphan identification is greater or equal than 1e-3 [17]. We made this threshold 1,000x more stringent and set it to >= 1.0 for all BLAST hits outside of the organism orphan belongs to. Each of the 100 taxa used for construction of query vectors had to be represented with at least 50 orphan proteins left after both stages of orphan filtering process. These proteins made the “stringent” orphan dataset. Probability of obtaining the same or greater number of taxonomic matches by chance, for each taxonomic rank can be calculated based on multinomial distribution. More formally, if we have r + 1 subsamples, *A*_*1*_, *A*_*2*_, …, *A*_*r*_, *A*_*r+1*_, then the probability of drawing at least *i*_*1*_, *i*_*2*_, …, *i*_*r*_ of elements *A*_*1*_, *A*_*2*_, …, *A*_*r*_ with replacement, in n repeated draws equals:

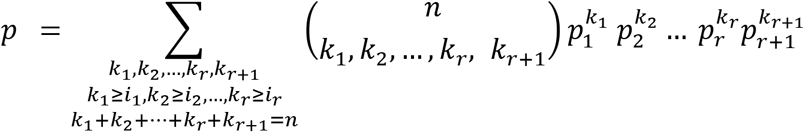

This formula considers the fact that dataset is imbalanced, meaning taxa distribution is not uniform. Due to our limited computational resources we could not calculate probabilities for taxonomic ranks that had more diverse composition, therefore we have calculated probabilities of a more probable event. This event can be described as drawing all positively identified taxa at least once. The same multinomial distribution applies, and if we have r + 1 subsamples, *A*_*1*_, *A*_*2*_, …, *A*_*r*_, *A*_*r+1*_, then the probability of drawing at least one of elements *A*_*1*_, *A*_*2*_, …, *A*_*r*_ with replacement, in n repeated draws can be described as:

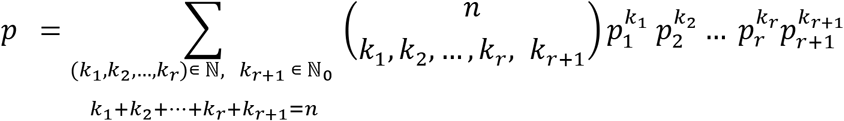

### Latent Taxonomic Signatures visualized

To produce illustrations of newly discovered shared sequence feature – Latent Taxonomic Signatures (LTSs), Savitzky-Golay filter was utilized [43]. This filter first convolves data by fitting successive sub-sets of adjacent data points with a low-degree polynomial by the method of linear least squares. Python includes this filter in Scipy (savgol_filter) so we implemented it within our code to generate plots showing LTSs of several selected organisms. For these plots, 3-peptide frequency data used for LSA species model training was used, smoothed by Savitzky-Golay windows length 999, and 4th order polynomial.

### Context dependent evolution

In order to make this experiment, we collected protein sequences from those Pfam families, which could be decomposed taxonomically into groups having at least 100 protein sequences per taxa. There were 468 such families fulfilling this condition on the species level. We collected an additional dataset consisting of protein family sequences grouped under genus level. Genus level dataset contained 201 Pfam families, and it was constructed to be independent from the species level dataset, meaning that it did not include proteins from taxa already included in the species dataset. Together, these two datasets shared 188 Pfam families; therefore difference in the genus dataset was 13 additional families. Species dataset aggregated protein sequences from 2,350 taxa, while the genus dataset aggregated protein sequences from additional 570 taxa, not included in species dataset. In total, 292,000 protein sequences (randomly sampled 100 proteins per taxa) and 2,920 taxa were part of this experiment. In order to produce taxonomically unbiased control samples, to serve as positive control for actual proteins encoded by genes of different organisms, we have used the Pfam protein family HMM models in order to generate 100 protein sequences per taxon, from a fully configured HMMER protein family search profile. This was accomplished using “hmmemit -p” option which is available in new HMMER3 program suite installed locally on our machine. We did this in order to obtain sequences fully consistent with a sequence family consensus, thus representing taxonomically unbiased, e.g. “selfish” behavior. This option allowed us to sample sequences from a fully configured HMMER search profile, so we could obtain homologous sequences by HMMER implemented HMM definition, including non-homologous flanking sequences, allowing us to come as close as possible to the theoretical template of the protein family. After collecting and sorting all protein sequences, taxonomically grouped Pfam family proteins were folded into LSA 400-dimensional vector space. Intra-class comparisons refer to comparisons between taxa vectors belonging to the same protein family. On the other hand, inter-class comparisons correspond to comparisons between vectors made from protein family proteins and Latent Taxonomic Signatures extrapolated from entire proteomes.

## Acknowledgments

We thank all those who keep contributing by making biological sequence data publicly available. We also thank prof. John Cullum, prof. emeritus Daslav Hranueli and prof. Walt Dunlap for discussions and constant encouragement. We thank late prof. Gabriel Dover for his work on molecular drive and concerted evolution. Without his scientific courage and vision, we would have great difficulties interpreting results herein presented.

This work was supported by Croatian Science Foundation (grant number IP-2016-06-3509) and Scientific Centre of Excellence for Marine Bioprospecting – BioProCro (grant number KK.01.1.1.01).

## Author contributions

E.M. and T.C. equally contributed to formal mathematics sections. J.D. contributed by developing web based search retrieval system based on LTSs, which can be readily accessed at http://matrix.pbf.hr/, J.Z. and P.F.L contributed to the writing of the manuscript and performed major revisions, A.S. conceived the manuscript, wrote the code and performed experiments. All authors analyzed the results and reviewed the manuscript.

## Data and materials availability

All of the datasets generated and analyzed during this study are available from the corresponding author (A.S.) on reasonable request with no restrictions since their source is already referenced and publicly available. All computer code generated during this study is also available from the corresponding author (A.S.) on reasonable request and from a GitHub repository: https://github.com/astarsky2016/Latent-Semantic-Signatures. Web application based on LSA taxa model is freely accessible at: http://matrix.pbf.hr/

## Supplementary Materials

Supplementary Table 1 – Information on taxa included in LSA species model

Supplementary Figure 1 – Protein tokenization scheme, cosine similarity and taxonomy assignation in voting scenario

Supplementary Table 2 – Table containing download links for FASTA files with “train” and “test” protein sequence sets used.

Supplementary Table 3 – Percentage of initial taxa query space as defined by available lineage data – used in SBH and VSM method-benchmarking tests

Supplementary Table 4 – Relaxed orphan sequence dataset

Supplementary Dataset 1 – Download link for multi-FASTA formatted sequences comprising “stringent” orphan dataset

Supplementary Table 5 – Protein family taxonomy based groups used in “selfish” vs “altruistic” mode of evolution experiment

